# Water as a reactant in the first step of triosephosphate isomerase catalysis

**DOI:** 10.1101/2021.01.30.427993

**Authors:** Max Yates, Patrik R. Callis

**Author notes:** **Corresponding Author** Patrik R. Callis-- to whom correspondence should be addressed. *Department of Chemistry and Biochemistry, Montana State University* Bozeman, Montana, 59717. United States., **Author**, Max Yates-- *Department of Chemistry and Biochemistry, Montana State University* Bozeman, Montana, 59717, United States.

## Abstract

The enzyme triosephosphate isomerase (TIM) performs a crucial role in the extraction of energy from glucose, doing so by converting dihydroxyacetone phosphate (DHAP) into glyceraldehyde phosphate, thereby doubling the yield of ATP molecules during glycolysis. The initial step of the mechanism is the seemingly unlikely abstraction of the *pro-R* methylene hydrogen from C1 by a conserved glutamate (Glu165), an assignment that has been both universally accepted yet a much-studied phenomenon for decades. In this work we introduce an alternative mechanism in which water as a strong general base abstracts the carbon proton acting effectively as hydroxide. We posit that strong electric fields associated with the substrate phosphate promote facile autoionization of water trapped near the phosphate dianion of DHAP and Glu165, an example of substrate assisted catalysis. Classical molecular dynamics simulations assert that the closest water oxygen atom is consistently closer to the *pro-R* H than the carboxylate oxygen atoms of the accepted base Glu165. Our proposal is further supported by quantum computations that confirm the implausibility of abstraction of the methylene hydrogen by glutamate and the ease with which it is abstracted by hydroxide. The necessity of Glu165 for efficient catalysis is attributed to its crucial involvement in trapping the vital water in an environment of high electric fields which promote ionization far more rapidly than in bulk solvent.

## Introduction

A most thoroughly investigated enzyme mechanism is that of triosephosphate isomerase (TIM), which by a simple isomerization of dihydroxyacetone phosphate (DHAP) to glyceraldehyde 3-phosphate (GAP) involving two proton transfers, doubles the energy yield from the glycolysis of glucose.^1^ The undisputed first step of this simple transformation, presented in Fig. S1, is the chemically unlikely abstraction of the *pro-R* methylene proton (green in Fig. 1) from the hydroxy methyl by the weak base Glu165, an edible amino acid commonly known as “mono sodium glutamate” or MSG. Despite universal acceptance, a few dozen papers published in the last 35 years have been driven by the goal of comprehending the environmental details within the enzyme that lead to such a phenomenal enhancement of basicity.^2–7^ To our knowledge, none have previously contested the validity of Glu165 as the catalytic base.

**Fig. 1.**
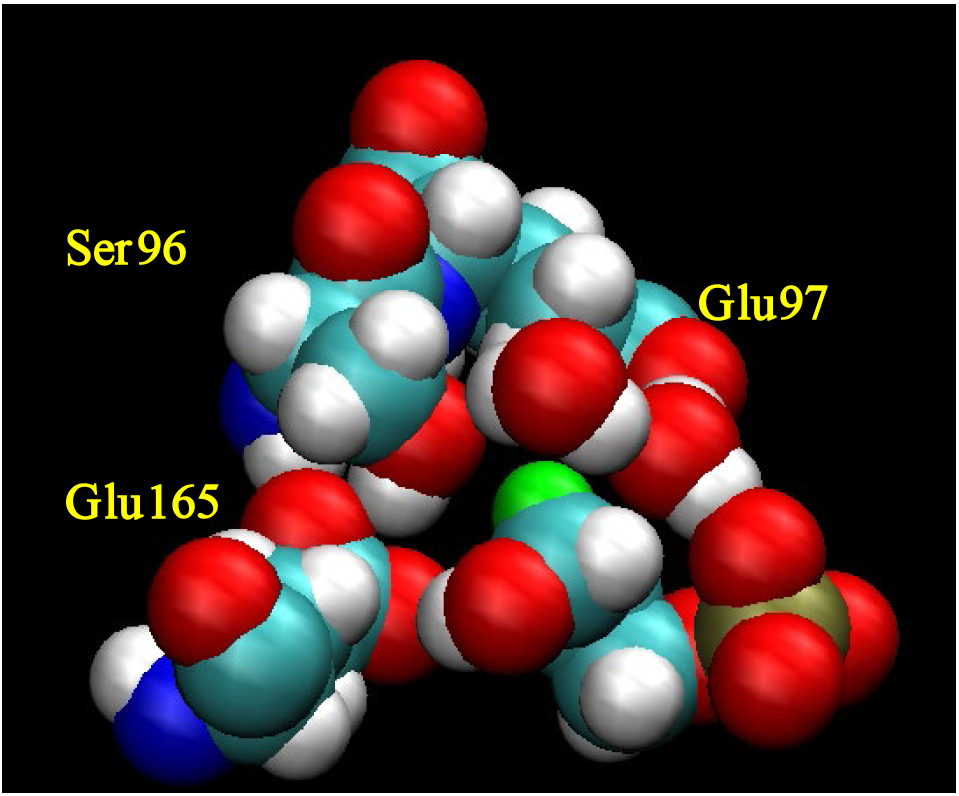
Snapshot from MD simulation of TIM, showing a commonly found “water wire” bridging from an O on phosphate P (olive at right) to OG of Ser96 (left). Ser 96 is shown making a strong bidentate grip on Glu165. The abstracted pro-R proton is in green.

The enduring acceptance has roots in the meticulously thorough isotopic study by Knowles and Albery published in eight back-to-back papers^8^, which unquestionably established that the *pro-R* proton on C1 appears in the solvent during catalysis--but never in the absence of enzyme. In a later review, Knowles^9^ articulated well-reasoned circumstantial evidence for the remarkable fact that the weak base Glu165 is able to abstract the very weakly acidic methylene proton. Four reasons were put forth: (1) the close proximity; (2) the bidentate glutamate after abstraction is immediately poised to protonate C2 of the enediol, as is needed for the isomerization; (3) the abstracted proton lies orthogonal to the enolate plane; and (4) the abstracting orbital of the carboxylate is *syn*, which has been argued to be much more basic than the *anti* orbital.^10^ A fifth point is made by Knowles that mutating Glu165 to alanine virtually eliminates the catalytic power of the enzyme.

The focus of this paper is to present circumstantial evidence that supports an alternative, chemically more credible mechanism wherein the catalytic base is *not* Glu165 but rather the strong base hydroxide, produced via proton transfer from a proximate water to the alkyl phosphate dianion of the substrate DHAP, which is a stronger base (pK_a_=7) than glutamate in water. We support this assertion by three types of computational observations, all motivated by the abundant access of water to phosphate and the *pro-R* proton.

1. Quantum computations that indicate difficulty of abstraction by Glu165 and show rapid abstraction by hydroxide.
2. Classical molecular dynamics (MD) simulations that: (a) consistently show trapped water oxygen atoms being closer to the *pro-R* proton than the oxygen atoms of the putative base Glu165; (b) elucidate the retention of measurable catalytic activity for the E165A mutant through reduced—but not abolished—proximity of water near the *pro-R* proton in the absence of Glu165.
3. Electric field calculations on the O-H bonds of water in the active site that consistently show higher fields that would promote ionization compared to those elsewhere in and around the protein, and greatly exceed those on the *pro-R* proton.

## Results

### Quantum computations of pro-R H abstraction

The basic question as to whether a glutamate is a sufficiently strong base to abstract a methylene proton can in principle be answered with a series of high-level quantum chemical computations. We have compared ab initio quantum molecular dynamic simulations using Atom-centered Density Matrix Propagation (ADMP)^11^ on a variety of small molecular clusters containing the pertinent DHAP, Glu165, and nearby water to those with the putative hydroxide using a range of methods/basis sets. In all cases these have given the result that Glu165 does not abstract the *pro-R* proton from DHAP, but that a hydroxide placed at the close positions found for water in the MD simulations abstracts this proton within 0.1 ps in a variety of model systems. (Movie S1) Ab initio optimization calculations confirm the activationless nature of the abstraction by hydroxide at H-O distances up to 2.8 Å. (Movie S2)

Additionally, quantum simulations were carried out on clusters more representative of the active site containing the essential active site residues of TIM, Lys12, His95, Ser96, Glu97, and Glu165 as well as water and DHAP, similar to what is shown in Fig. 1 (where Lys12 and His95 are omitted for clarity). Nearly 300 independent simulations from various conformations of the protein taken from the MD simulations were performed with various basis sets. In none of these runs was the *pro*-*R* C1-H bondlength seen to be lengthened significantly towards Glu165.

### MD simulations implicate active site water

Classical molecular dynamics (MD) simulations of TIM solvated in explicit SPC/E water consistently show much water in contact with the active site ensemble. During the majority of the over 250 ns of simulation shown in Fig. 2, a hydrogen-bonded network of water exists, stretching from the phosphate towards Glu165 and Ser96. Fig. 1 shows such a network consisting of a 2-water molecule “wire” bridging between the phosphate in which the O of the water H-bonded to the O of Ser96 is effectively in contact with the *pro-R* proton (green) of C1. Such a water is commonly found throughout the simulation and is often H-bonded to the OG of the conserved residue Ser96, a residue long recognized for controlling water near in the active site.^12^ Fig. 2*A* compares the distances from the OE atoms of Glu165 (red and green traces) with the distance of the water O atom closest to the *pro-R* H (blue). The plots show that the water molecule closest to *pro-R* H is generally slightly closer than the closest carboxylate O atom of the putative catalytic base Glu165. The same data is shown as histograms in Fig. S3. These simulations were carried out with the phosphate in the di-anion protonation state, whose conjugate acid has a pK_a_ of 7.1 in pure water. Therefore, the dianion form is expected to exist about 50% of the time in neutral aqueous solution. The environment in the active site could shift the equilibrium to being predominately in the dianion form.^13^ Fig. 2*A* also reveals the effect of fluctuations of enzyme conformation. These are typically caused by changes in positions of Ser 96 and Glu165 relative to the *pro-R* proton. Details regarding the large-amplitude changes are given in SI. In the context of the previous paragraph, we note that another reason for the uncontested acceptance of Glu165 as the cata-lytic base is that the E165A mutant is nearly 10000 times less effective in terms of rate^14^. This means, however, that catalytic activity of this mutant is not completely abolished despite the absence of a catalytic base in a position to abstract the C1 proton. We therefore performed a 50-ns MD simulation on the E165A mutant form and compared the distance of closest water with the wild type distances in Fig. 3. We find that for E165A, a network of water linked to the phosphate is retained that places water near *pro-R*, but the average distance of the water O atom is 3.3 Å, which is considerably longer than the average distance of 2.7 Å for the wild type. Perhaps more significant is the paucity of large fluctuations for E165A. The minimum distance to OE is 2.8 Å whereas there are many distances < 2.3 Å for wild type.

**Fig. 2.**
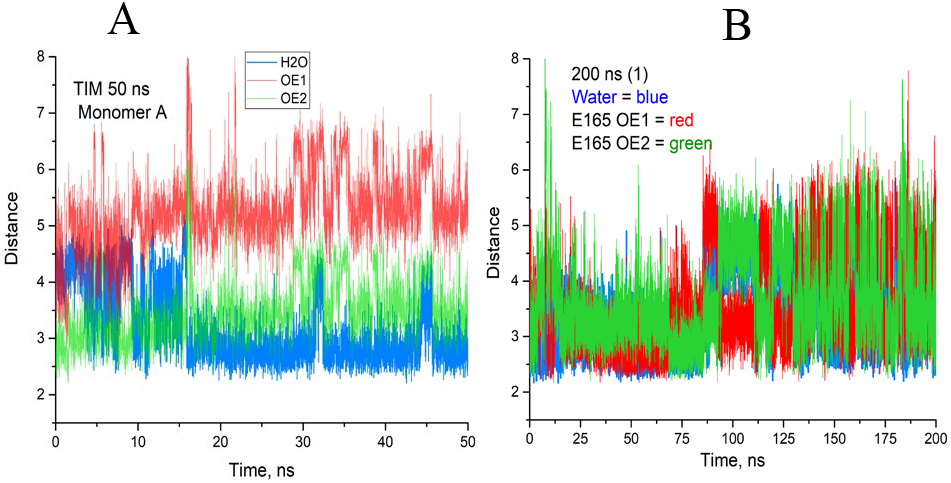
plots of Glu165 OE (red and blue) and the closest water O distances to the pro-R H (blue) on C1 of TIM

**Fig. 3.**
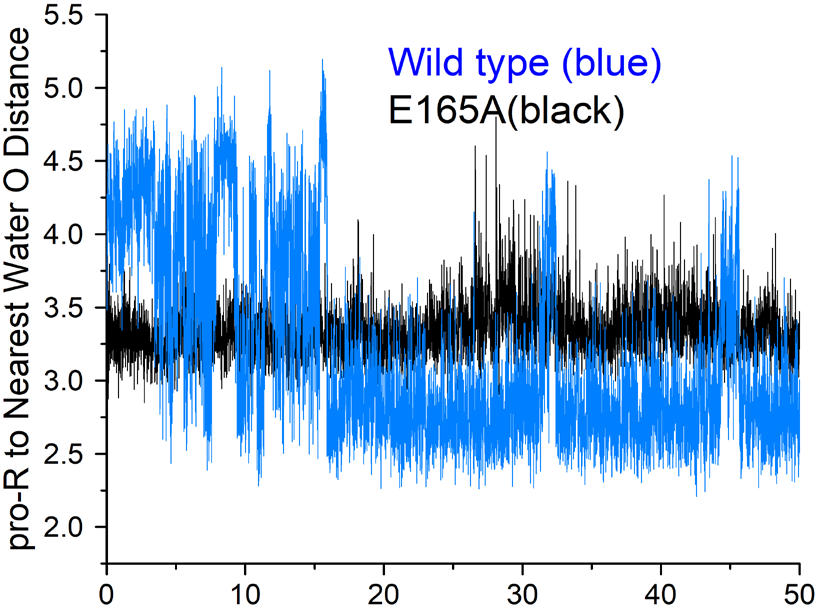
Plots of the closest water O distances to the pro-R H on C1 for wild type TIM (blue) and of the closest water O distances to the pro-R H on C1 for E165A of TIM.

Overall, these findings are consistent with the observed^14^ retention of weak enzymatic action for E165A, for which the primary cause of the longer *pro-R*-water distances could be that, without Glu165, the OG of Ser96 is not held tightly in a position to hold a water molecule near the *pro-R* proton. The importance of Ser96 regarding water has been stressed^12^, and noted in the context of fully appreciating shifts in positions of water for drawing conclusions from mutations of active site residues.^15^

### Electric field computations

The work presented here is part of a larger effort directed to understanding enzyme mechanisms in general, the impetus for which comes from calculations of local electric fields in proteins performed in this laboratory for the purpose of attaining a molecular-level understanding of the very useful experimental variations of tryptophan fluorescence wavelength and efficiency in proteins.^16^ Our quantum calculations have convincingly demonstrated that fields on the order of 50 MV/cm (0.01 au) are required to attain the observed variations. Here we report results of investigating the strength of electric fields exerted on water in the active site of TIM using techniques similar to those of several groups who have extracted useful correlations of local electric fields and IR spectra of a variety of systems using classical MD simulations calibrated by high level quantum computations. These have been found to be extremely effective for understanding the IR and Raman spectra of pure water and various carbonyl probes embedded in proteins.^17–22^.

Results from the present work are shown in Fig. 4. In Fig. 4*A* two gaussian distributions for the electric fields on protons within 3.0 Å of phosphate oxygens are seen: one exhibits fields close to those of bulk water and the other exhibits a maximum at fields nearly double that for bulk water. The latter corresponds mostly with the protons H-bonded directly to the phosphate O atoms and the former to the other proton on those water molecules. We have seen very similar results for a collection of 40 GTPases and 30 ATPases (Figs. S4 and S5) In contrast, electric fields on the *pro-R* and *pro-S* protons are miniscule in comparison, as seen in Fig. 4*B*, whereas the fields on the protons of the water closest to the *pro-R* proton have a sizeable distribution with high fields.

**Fig. 4.**
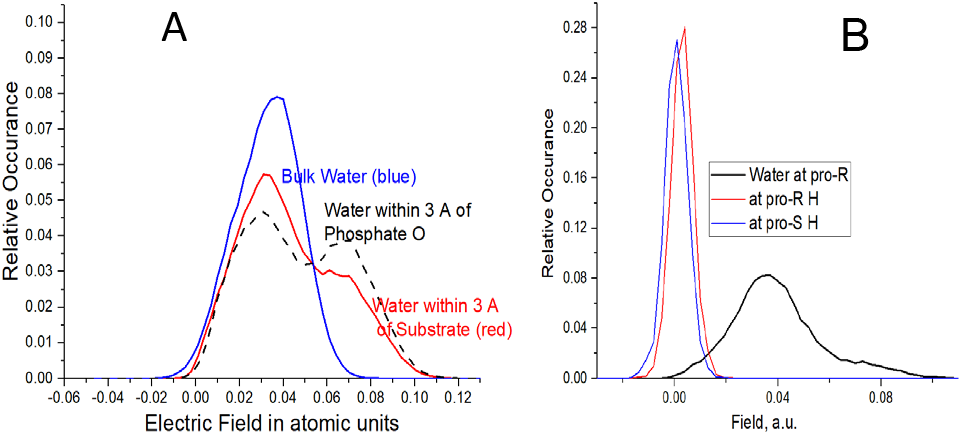
Histogram of computed electric fields along projected along the O-H bonds of bulk solvent water (blue), water within 3 A of DHAP (red), and water within 3 A of the phosphate O atoms (black) (*A*), fields for the closest water to *pro-R* H (black), the fields at the *pro-R* H(red) and *pro*-S H (blue) (*B*).

A connection between proton transfer in pure water and local electric fields from the water dipoles was made in ground-breaking papers concerning the precise mechanism of the autoionization of pure liquid water,^23–26^ indicating that these rare events occur only during solvent conformations that produce fields of 0.050-0.055 au on the transferring proton. In the present work we find that when a constant field of 0.055 au is applied to an *ab initio* energy-optimized water dimer, proton transfer to produce hydronium-hydroxide in vacuum occurs in 80 fs during an ADMP quantum MD computation using b3lyp/ccVTZ; a field of 0.06 au, applied to the same, causes a complete proton transfer within 15 fs. (Movie S2) Consistent with this finding, optimizing geometry starting from the same point with a constant applied field of 0.065 au shows barrier-less transfer. (Movie S3) Because part of the field on the transferred proton comes from the accepting O, we find that using the accepted point charge values assigned to SPC/E water for the purpose of MD simulations, a total field of ca. 0.09 au would cause hydroxide production. As can be seen from Fig. 4, for water in a position to be activated near the pro-R proton, fields exceeding 0.09 au are far more common than in bulk water, thereby adding to the plausibility that in the presence of phosphate dianion, hydroxide may be produced at a rate several orders of magnitude above that in pure water.

A schematic of our proposed alternative mechanism for the first step of the TIM mechanism is presented in Figs. 5 and S2. Here the catalytic base is a hydroxide produced by a concerted double proton transfer to the phosphate, although our MD simulations suggest that many variations on this theme are possible. The evidence presented above suggests that only hydroxide is a strong enough base for this abstraction. If this is the case, the enediolate ion would also be a strong base and could easily abstract a proton from water in the second step of the mechanism.

**Fig. 5.**
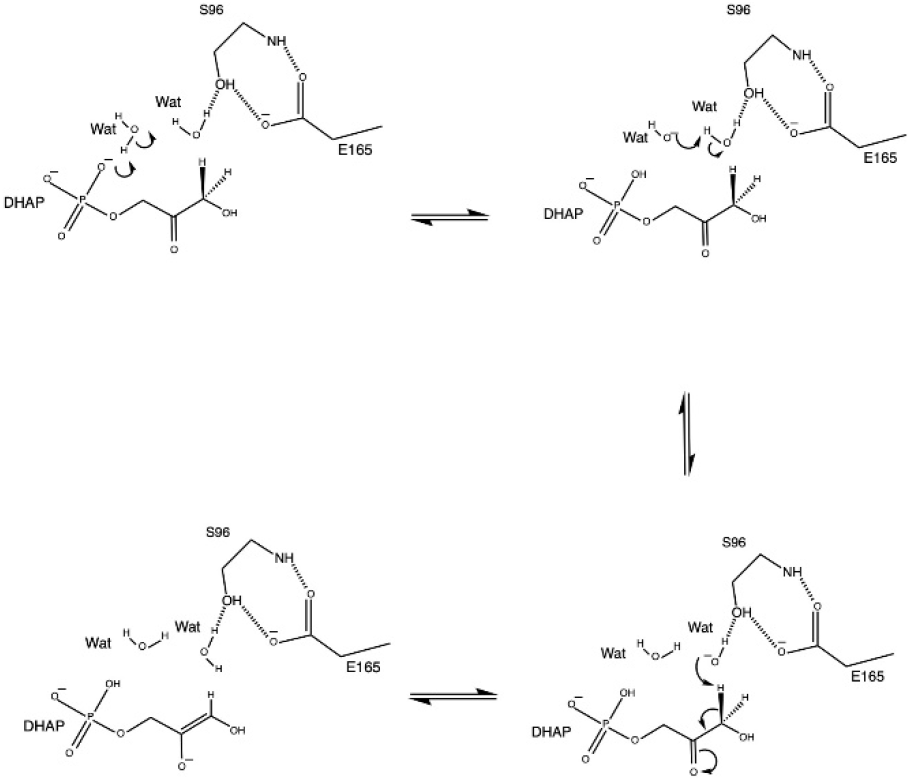
Proposed alternative pro-R abstraction scheme. Proton transfer from water to the phosphate dianion (possibly involving one or more water molecules) creates hydroxide, which rapidly abstracts the pro-R proton.

## Discussion

What is proposed above is an example of the concept of *substrate assisted catalysis*, which has been previously posited in the context of phospho-ester hydrolases and polymerases.^27–31^ Indeed, one of the earliest instances of such a proposal grew from skepticism regarding Asn61 as the general base in the mechanism of RAS p21 proteins^27, 29–31^, a motivation that appears parallel to that pursued here. More recently are reports involving *water* as the ultimate catalytic base stemming from water deprotonation involving phosphate^32, 33^, including computational studies that compare different detailed mechanisms, all involving proton transfer from water to phosphate at some point.^34, 35^ This concept does not seem to appear in previous work on TIM, however.

There is reticence in these works to use the word *hydroxide*. We are not necessarily proposing that a free hydroxide is produced during pro-R proton abstraction; the process could be concerted involving two or more water molecules—as in water autoionization and many other mechanisms involving water, and might better be described as water acting as a strong general base.

### QM

TIM has naturally attracted a number of high-quality quantum based computational studies.^36–39^ These uniformly have found values for activation barriers that are more or less in agreement with the barriers published by the extensive study of Albery and Knowles.^40^ Cui and Karplus^36^, found that virtually the entire effect of lowering by the active site was, not surprisingly, due to protonated Lys12, which is H-bonded to the O2 of the enediol anion. Xiang and Warshel, who applied frozen DFT, have also stated that they find good agreement for the proton transfer in TIM.^39^

The above cited works were carried out before the more accurate crystal structure with the true Michaelis complex was available.^41^ One of the few such computational studies using co-ordinates from that study was by Guallar et al^42^, who unlike previous work, and pertinent to the current work, found that having the phosphate protonated considerably lowered the barrier for abstraction of the *pro-R* proton. This would be expected, given that the proton is moving away from the negative charge on the phosphate.

More recently a comprehensive series of empirical valence-bond (EVB) investigations of activation free energies have also shown reasonable agreement with the experimental value^5, 43^.

### MD

The few crystallographic water molecules included in the ab initio computations of free energy barriers cited above are not representative of what is seen in our classical MD simulations. Crucially, there appears to be nearly complete access of water to Glu165 carboxylate atoms via water “wires” leading to bulk solvent water. This implies that the effective pK_a_ of Glu165 may be close to 4 as seen for aqueous solutions, in agreement with Hartman et al. who reported a value of 3.9 in the active site.^44^ (See SI for further information and discussion on pKa values in the enzyme)

This observation is not in accord with that of Kamerlin and co-workers^43^ who have recently presented comprehensive theoretical evidence for Glu165 as the catalytic base and who contend that only during transient periods, a water-excluding hydrophobic loop (loop 6) lowers the activation free energy and raises the pKa of Glu195 adequately for the abstraction of the proR carbon proton by Glu165. In our view, this popular view is somewhat circumstantial—not unlike that presented here.

The different protein conformations under which catalysis appears likely to proceed through a hydroxide-like intermediate according to our MD simulations may only be the “tip of the iceberg”. A larger number of independent simulations will be required to learn whether this is true.

### Electric fields

The subject of strong electric fields is important here because of arguments that glutamate is not a sufficiently strong base to abstract a carbon proton, and therefore requires extraordinary enhancement from the enzyme environment. One likely environmental effect provided by the enzyme would be extreme electric fields arising from the configuration of the several charged residues comprising the active site. It is evident from Fig. 4*A* that ~40% of active site water near the phosphate experiences an average electric field exceeding double the average found for bulk water. By the same methods, we find an order of magnitude smaller fields tugging on the *pro-R* and *pro-S* protons on C1 as seen in Fig. 4*B*. One might well ask how this can be, given that the OE atoms of Glu165 are often positioned to provide strong electrostatic attraction. The answer emerging from our calculations is that indeed the OE atoms do apply fields approaching the highest seen on phosphate water, but equally strong local fields produced by phosphate and the O atoms of C1 and C2 negate those fields.

The importance here of high fields is underscored by increasingly strong evidence based on vibrational Stark effects in the active site of delta ketosteroid isomerase (KSI) ^22^ and from seminal work on the autoionization of water.^23, 25, 26^ These studies stressed the importance of high electric fields in “pre-organization” that stabilizes the reaction transition state and both echo one of the primary principles of enzyme proficiency, first articulated by Warshel and Levitt in 1978.^45^ In autoionization, the organized element is essentially a donor-rich water molecule separated by a wire of compressed H-bonds involving two waters, which was deduced by examining the solvation pattern observed around the neutral water that had been a hydroxide immediately before the neutralization events.^25, 26^ Fig. 1 shows a commonly observed arrangement of water from our MD simulations wherein the phosphate takes the position of hydroxide, and one can easily visualize fluctuations that could compress the intervening H-bonds simultaneously, thereby facilitating a concerted, 2-proton jump leaving a hydroxide in position to abstract the *pro-R* proton. Pertinent here is that the simulations revealing details of water autoionization suggest that the rate of proton transfer is much higher than the net production of ions that leads to pH 7. This is because of rapid geminate recombination of the ions before separating,^23, 25^, which may not be so likely to happen within electrostatic confines of the enzyme.

### A hidden contradiction

A perplexing aspect of the literature concerning the requirement of extremely enhanced basicity of Glu165 in TIM is revealed in our classical MD simulations of Fig. 2. Namely, Glu165 is almost never without a hydrogen bond to at least one water molecule and/or the O1 hydroxyl, both of which are far stronger acids than a methylene C-H. This raises the question as to why this well-accepted “super-base” would not always remove a proton from water or from the O1 hydroxyl, and therefore virtually never be free to act as a base. A corollary of the super-base assumption is that during the instant when Glu165 is not protonated, a hydroxide would have been produced, which quantum simulations assure us will instantly abstract a carbon hydrogen.

The apparently successful quantum computations of activation barriers cited in the QM section above notwithstanding, the lack of a satisfying mechanism has persisted over two decades of searching for how glutamate becomes such a strong base. Occam’s razor would point to a simpler explanation: that the enzyme is *not* increasing the pK_a_ of Glu165 enormously, but rather, hydroxide produced by the more basic phosphate dianion is the most reasonable catalytic base. This conclusion is entirely consistent with studies focused on the ability of added free phosphate and phosphite to restore enzymatic activity in TIM when bound by a truncated form of DHAP lacking the phosphate group.^46^ The absence of proton transfer to Glu165 in our ADMP quantum dynamics simulations, of course, does not rule out such a transfer because the total simulation time was only ~1 ps, which is not in the realm of the measured kcat for proton abstraction time of milliseconds.^40^

### Related carbon proton abstractions

The findings presented above apply to other isomerases involving stereospecific abstraction of a carbon proton by weak, acetate-like catalytic bases.^19,47, 48^ Most of these involve phosphorylated sugars except for KSI, for which the abstracted proton is reported to have a pK_a_ of 137, suggesting that abstraction by a carboxylic base in KSI is not unreasonable.

## Conclusions

We have presented evidence supporting an alternative to the unproven—yet uncontested—greatly enhanced basicity of Glu165 as the catalytic base in the initial step of the TIM mechanism. We conclude that an equally compelling explanation is that hydroxide from water ionized by the DHAP phosphate di-anion is the catalytic base, i.e., the usual suspect in most enzymatic reactions requiring extreme measures. The conserved and virtually essential Glu165 is here reassigned to the role for which it is apparently uniquely qualified: acting in concert with conserved Ser96 to supply a necessary role of trapping water in a manner such that a hydroxide can often be found adjacent to the *pro-R* methylene hydrogen on C1.

This conclusion emerged almost seamlessly from our decade-long study of the effects of strong electric fields from protein and water on tryptophan fluorescence wavelength and brightness. A constant theme of those studies was the necessity for including explicit bulk water whose collective field made essential contributions.^16, 49^ In the present study, the contribution of electric fields was strongly reinforced by the observations from the literature pointing to such fields being implicated in the autoionization process that provides the well-known pH 7 of liquid water.

We believe this conclusion is more universally applicable than is presently appreciated. The possibility exists that many enzymes must efficiently dissociate water into hydroxide and hydronium ions to achieve the observed catalytic prowess. That is to say, the extreme electrostatic environment of enzyme active sites can make *water* act as a “super acid/base”. If true, one may say that water is an essential cofactor for such enzymes.^50^

## ASSOCIATED CONTENT

### Supporting Information

Computational methods, details of the origins of large fluctuations seen in Figs. 2 and 3, further elaboration on the pertinent question of enzyme-pKa of Glu165 and phosphate, Figures S1-S6, and Legends for Movies S1, S2, and S3.

## ACKNOWLEDGMENT

The authors thank Drs. James Vivian, Pedro Muino, Matt Cook, John Kiely, J. Michael Schurr, and John Straub for helpful conversations.

---

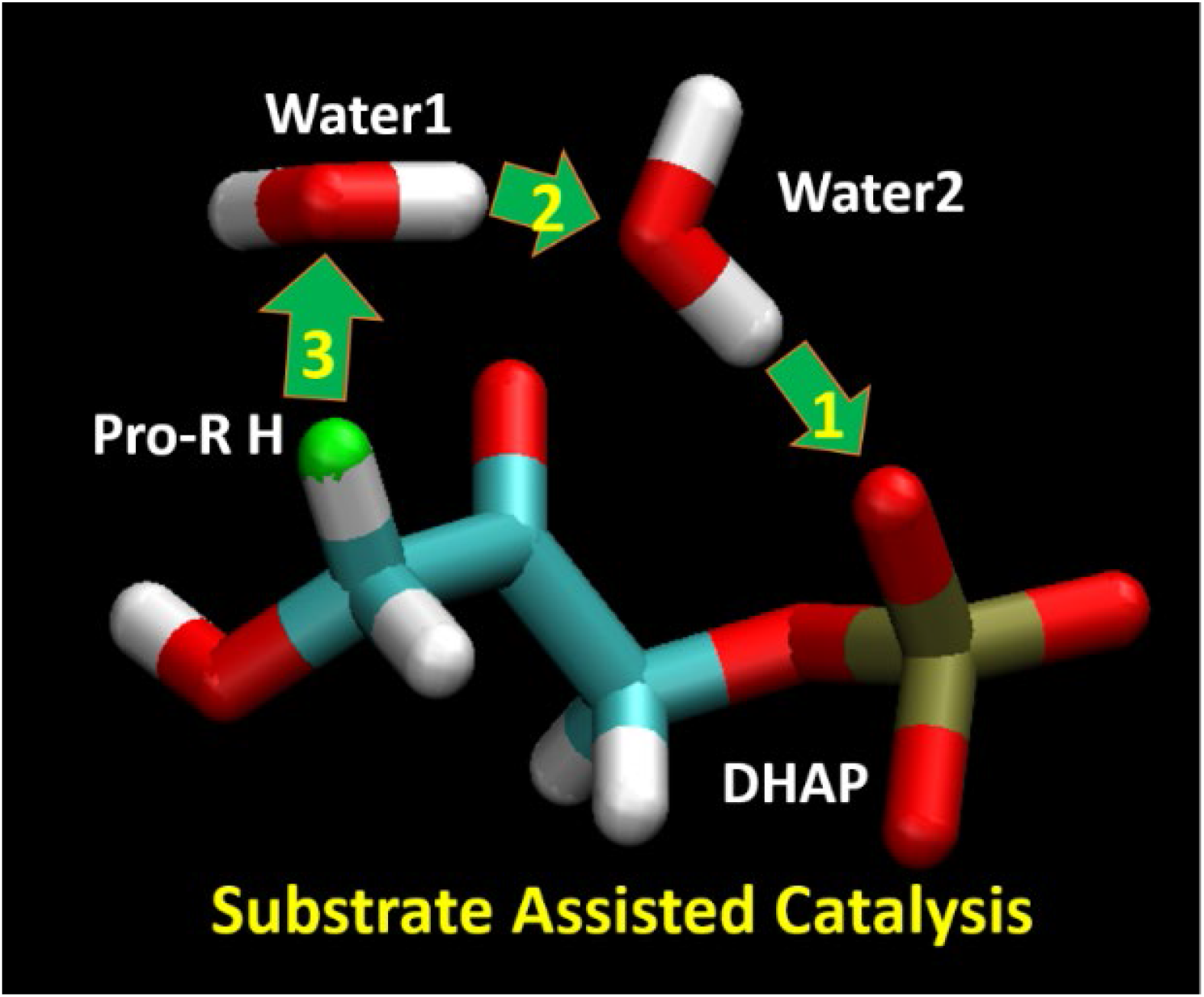
TOC graphic.

