## Supplementary material for "Water as a reactant in the first step of triosephosphate isomerase catalysis": Figures and Movies

Figure S2. . ....................................................................................................................................... 3

Figure S3. . . ..................................................................................................................................... 3

Figure S4. . . .....................................................................................................................................4

Figure S5. . . .....................................................................................................................................4

Figure S6. . . .....................................................................................................................................5

Movie S1. . ...................................................................................................................................... 6

Movie S2. . ...................................................................................................................................... 6

Movie S3. . ...................................................................................................................................... 6

##### Materials and Methods

Quantum MD used the method called Atom-centered Density Matrix Propagation method provided by Gaussian09[1] using the keyword ADMP.[2,3] This method describes quantum zero-point and tunneling effects, unlike classical MD, and is stated to provide results equivalent to those from Car-Paranello [4].

Gaussian basis functions instead of primarily plane waves. Energy optimizations were also carried out with Gaussian09, with HF/3-21, b3lyp/6-31g(d,p), and b3lyp/cc-pVTZ.

Classical molecular dynamics were performed with Gromacs version 2016.3 [5] using the SDSC Comet resources. The initial model of the reactant bound state was taken from a high-resolution structure of TIM generated from x-ray crystallography (RCSB PDB, 1NEY) [6]. The inhibitor molecule was modified to the substrate DHAP, which had been manually added to the topology database. The structure was solvated in a box with a volume of 772.11 $nm^3$ using the SPC/E water model. Simulations used periodic boundary conditions and achieved energy minimization through the steepest descent algorithm. The Amber ff99SB -ILDN force field [7] was used in all simulations, which was performed independently for 10, 50, and 200 nanoseconds.

Electric field calculations employed a homegrown Fortran77 procedure that independently very closely duplicated results published by Skinner and co-workers.[8,9] The input for the program are files in PDB format extracted from the .xtc MD trajectory files generated using Gromacs as noted above.

**Causes of distance fluctuations seen in Fig. 1*(A)* of text.** During the early 15 ns of the simulation, there is little close water, which is a period during which Ser96 is not yet in contact with Glu165 and water is essentially excluded. Thereafter, however, ~10-15 ns-periods of uninterrupted water trapped near the proton to be abstracted are common. The large transient distance increases from OE1 and OE2, at 29 and 32 ns, for example, correlate strongly with the O1-C1-C2-O2 dihedral angle, which in turn correlates strongly with whether HO1 is H-bonded to Glu165 OE2, Ser96 OG, or to a water h-bonded to Ser96 OG, i.e., internal torsional motions within DHAP. Transient regions of the trajectory for which the closest water makes sudden increases to near 4 Å correlate well with the intrusion of the large hydrophobic side chain of Ile170, which is effectively in van der Waals contact with the *pro-R* proton during such intervals.

In addition, during the exceptional upward fluctuation in the OE distances at 16 ns, DHAP is seen to make a short separation from Glu165. The closest water at the same instant becomes much closer by virtue of an instantly constructed 3-water wire stretching from phosphate to Ser96 OG, which remains attached to Glu165 by the double H-bond backbone "grip", which in turn allows DHAP to regain its H-bond to Glu165.

**Causes of distance fluctuations seen in Fig. 1*(B)* of text**). During the 200 ns run MD simulation DHAP is initially H-bonded to Glu165 OE2 via HO1 with a very short proR-OE2 distance and a close water, which remarkably remains trapped between OE2, Leu230 O and Gly210 for 54 ns, with another water bridging to phosphate. From 54-64.5 ns it alone bridges from OE2 to phosphate. The large spike to ~8 Å for the OE2 distance at from 7.4-10.5 ns coincides with DHAP becoming detached from Glu165 and re-attaching at 8.5 ns. During the 0-10.5 ns, Ser96 has no significant grip on Glu165. Histograms of the distances plotted in Fig. 2*B* are shown in Fig. S2.

**Further discussion of $pK_a$ in the enzyme.** Further circumstantial evidence comes from the TIM enzymatic activity vs. pH study of Plaut and Knowles [9] in which maximal activity was observed between pH 6 and 8.5, and fell off steeply outside those limits. This is not consistent with the normal $pK_a$ of glutamate in proteins, but Glu165 was considered a reasonable candidate for the titrating base, even though the $pK_a$ for the glutamate side chain is typically given as ~4. The Plaut-Knowles pH study is ambiguous because the $pK_a$ of Glu and Asp can be as high as 7 when buried in a protein, while the normal $pK_a$ for DHAP is 6, entirely due to the phosphate. As noted in the text, however, the abundant access of water to Glu165 carboxylate, suggests that environment should *not* be considered "hydrophobic". Plaut and Knowles also used the argument from Eigen[11] that all common bases, with oxygen, nitrogen or sulfur centers, combine with protons in aqueous solution with a second-order rate constant around $10^{10}$ $M^{-1}$ $s^{-1}$, concluding that "this requires that an enzymic acid group must dissociate with a rate of 20000 $s^{-1}$ and must have a pKa < 6, unless proton tunneling is invoked".

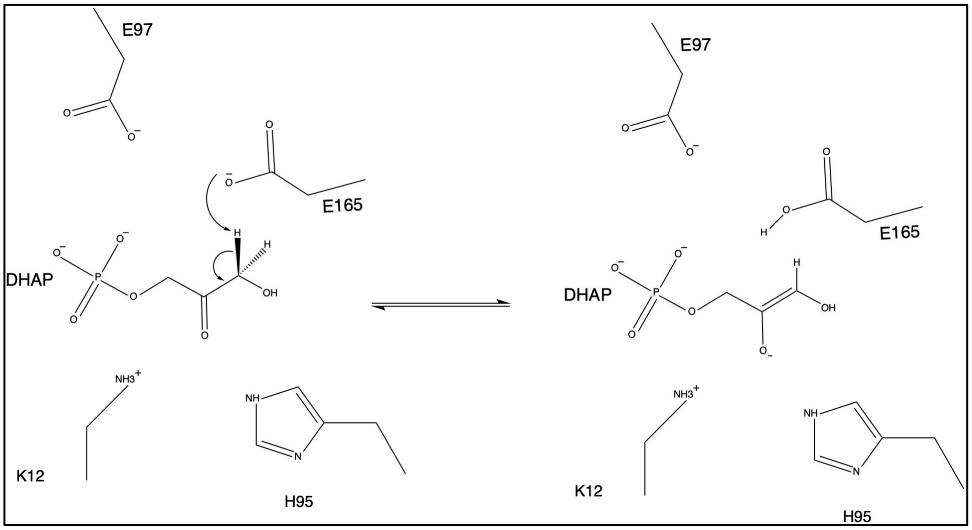

**Fig. S1.** The long-accepted mechanism of *pro-R* proton abstraction from $C_1$. The E165 anion is the base that abstracts the proton. The other active-site members shown are *not* involved in the initial step

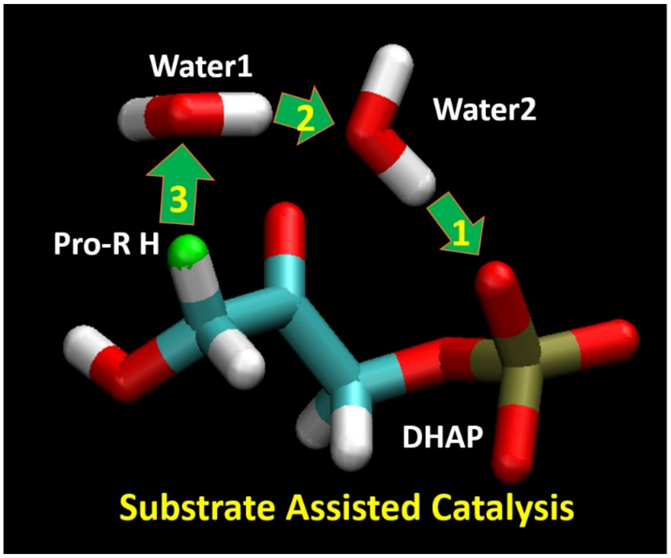

**Fig. S2**. A cartoon depicting the alternative mechanism proposed in Fig. 5 of the text, and is identical to the manuscript Table of Contents Figure.

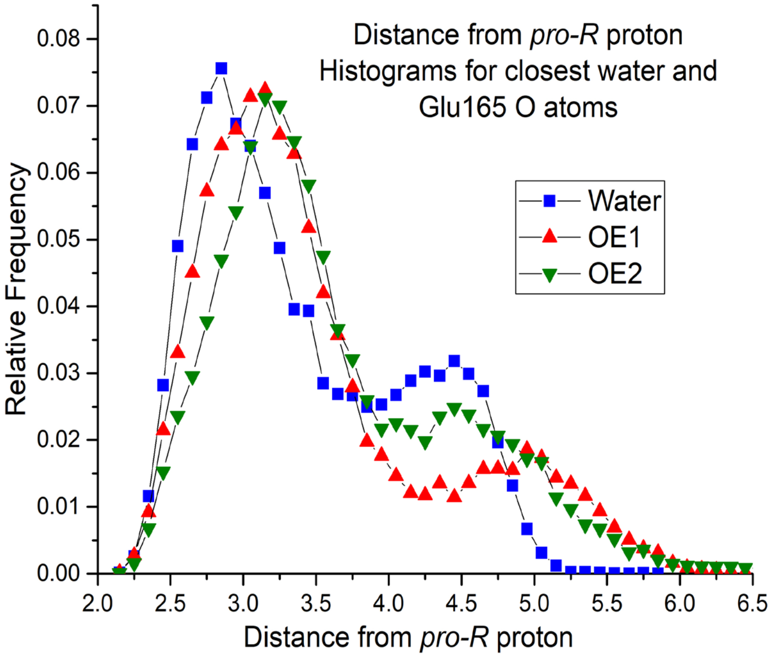

**Fig. S3**. Histograms of the distances found from the *pro-R* proton to the closest water oxygen, and to the OE oxygens of Glu165. Water in the active site is ubiquitous in MD simulations and is seen to be slightly more available for the formation of hydroxide proximal to the *pro-R* proton compared to OE distances.

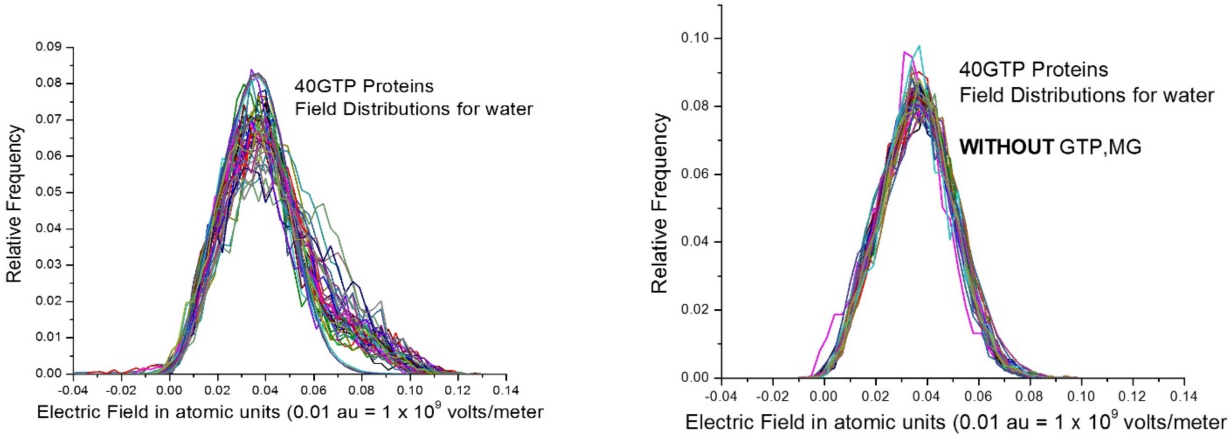

**Fig. S4**. Histograms of electric fields for **water in the active site** for 40 GTPases, **with and without GTP-Mg$^{2+}$**.

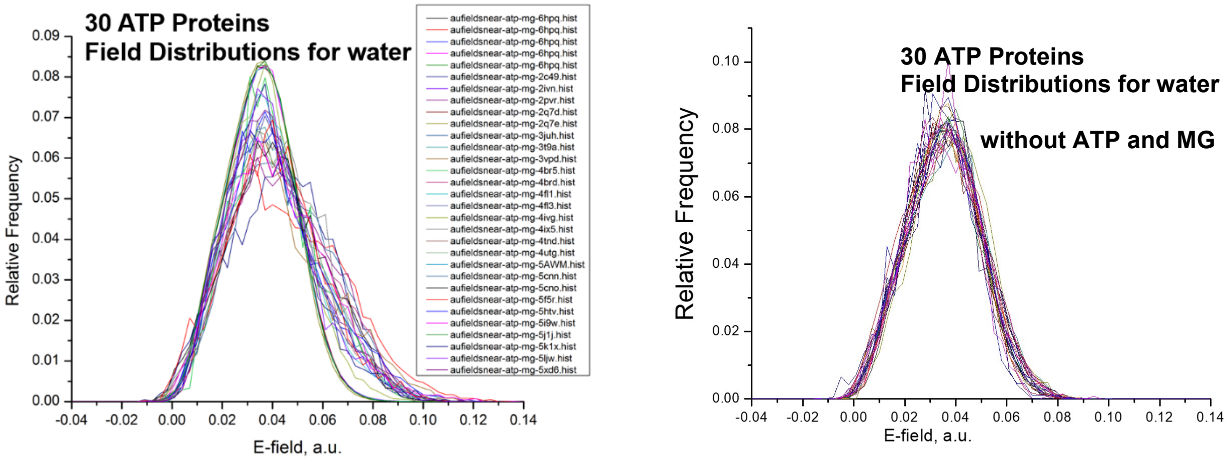

**Fig. S5**. Histograms of electric fields for **water in the active site** for 30 ATPases, **with and without ATP-Mg$^{2+}$**.

### Water as an Essential Cofactor for all Enzymes

Pedro L. Muíño[1] and Patrik R. Callis[2].
[1]Saint Francis University, Loretto, PA, USA, [2]Montana State University, Bozeman, MT, USA.

#### ABSTRACT

Enzymes are essential for the chemical reactions of all living organisms. Water is widely acknowledged as being important to enzyme action, but at the same time, a *common textbook attribute of enzymes is that they protect the active site from water*. We propose here that a more precise statement is that **the universal function of enzymes is to provide a pathway for water to the active site, where it is activated**. **We further propose that this is true for all enzymes**. This is to say that water is an essential co-factor for all enzymes, much like $Mg^{2+}$ ion is for ATPase action. We support this point of view with: (1) a survey of 55 crystal structures of enzymes arbitrarily chosen from all 6 classes of enzymes, all of which exhibit a "water wire" leading from bulk water to the active site; (2) a survey of electric fields in nucleic acid transferases and hydrolases along water O-H bonds (in the presence and absence of substrate) that generally reveals much higher fields for water trapped near enzyme active sites in the presence of substrate; (3) a survey of electric fields on the ribose O2'-H2' bonds that reveals a larger electric field on the nucleotides bond to be cleaved.

#### 1. Introduction

All biochemical reactions require one or more enzymes.

Enzymes enormously accelerate the rates of chemical reactions over the rates of the same reactions in water (by $10^{8}$-$10^{17}$-fold),

The precise manner by which enzymes accomplish this in detail is still considered an open question

This poster is about introducing a novel hypothesis stating that
(1)**intense internal electric fields** in proteins **profoundly affect the properties water in active sites**, and
(2) ALL enzymes incorporate water wires leading from bulk to their active site.

The opinions to be presented are in part motivated by experience with how these **same intense internal electric fields** have previously led to an **understanding of tryptophan fluorescence behavior**. For example, a field of $5 \cdot 10^{9}$ V/m leads to fluorescence quenching when oriented to force electron transfer from the indole ring to an amide backbone. The magnitude of this field is equivalent to 1000 times the field needed to cause an arc in air.

#### 2. Computational Methods

PDB files for enzymes crystal structures with a resolution < 1.5 Å were selected from the RCSB database. Water wires were detected via an algorithm that selected those water atoms within 4 Å of the active site amino acids. Then, selected water atoms within 4 Å of the first set of atoms and so on iteratively until either no new water atoms are selected or the wire has reached the protein surface

.

To calculate the electric fields on the pertinent O-H bonds, the following procedure was used:

- The PDB structure of the enzyme (plus substrate, if used, cofactors, water of crystallization) was used as input for molecular dynamics using GROMACS (AMBER99SB-ILDN forcefield) after including an explicit SPCE water field and, if necessary, parametrization of the substrate.
- Resulting structure was minimized and equilibrated for 200 ps to reach thermal equilibrium at 300K.
- Dynamics were then run for 10 ns, with coordinates extracted every 10 ps.
- A homemade code was used to calculate the projection of the electrostatic field onto each of the O−H bonds of bulk water and those in contact with the active site atoms.

#### 3. Water Wires

Water "wires" (yellow) within 4 Angstroms of the active site residues. The procedure described in the Computational Methods produced a water chain leading from the active site to the surface of the protein for **ALL 55 crystal structures examined.**

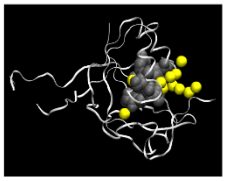

Human gamma-glutamylamine cyclotransferase complex with 5-oxoproline
(PDB code 3JUC)

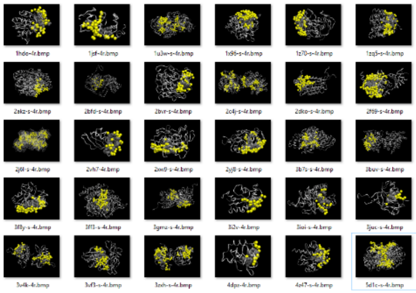

The composite to the left shows 30 of the 55 enzymes that we have analyzed.
The protein is shown as a ribbon, the atoms of the active site are shown explicitly in dark gray and the oxygen atoms of the water wires are shown in yellow.
**All 55 enzymes examined show water wires from the active site to the protein surface (i.e. the bulk)**

#### 4. Electric fields on the O−H bonds of water

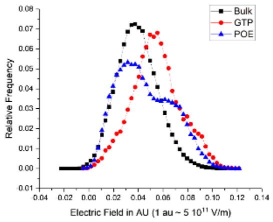

- The graph above for a bacterial polynucleotide kinase bound to GTP and RNA shows a histogram of the exceptionally high electric fields felt by the water molecules near the active site to a nearby phosphate compared to those for the O-H bonds in bulk water.
- The presence of the substrate is necessary to observe this enhanced field near the active site.
- Many enzymes (but not all in our calculations) showed this differential behavior.

#### 5. Electric Fields on Ribose O2'−H2' Bond

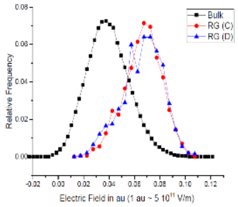

- The graph above (for a bacterial polynucleotide kinase bound to GTP and RNA) shows a histogram of the exceptionally high electric fields felt by the O2'−H2' bond of the RG nucleotide. The high field is the result of a tight H-bond with the O3' phosphate joining to RG to RC. While this is not the site of cleaving, we postulate that the O2' proton transfer in the proximity of an active site would almost certainly generate hydroxide that would be the strong base required for nucleophilic attack of the scissile bond.
- .

#### 6. Summary

1. Electric fields on water protons in the active sites of the few enzymes we have examined are often MUCH stronger than those of pure liquid water.
2. We believe the rate of hydroxide formation within active sites is therefore many orders of magnitude faster than in bulk water and may be a primary factor in the extreme catalytic power of enzymes.
3. We postulate that no enzyme functions without water, and therefore that a wire, or channel of water will be found for ALL enzymes.
4. Large electric fields are almost always found near the enzyme-substrate complex. Water buried elsewhere in the enzyme is typically indistinguishable from bulk water, even when in the presence of (non reacting) chemically identical structures.
5. These fields are specifically observed only at the site of reaction. If true: **WATER IS A COFACTOR**.

#### *Acknowledgment*

The authors thank the XSEDE Supercomputer center for access to computational time and resources

**Fig. S6**. Poster from Ref. 47: Muino, P. L.; Callis, P. R., "Water as an Essential Cofactor for All Enzymes". *Biophysical Journal* **2020,** *118* (3), 535A-535A. (Abstract only)

**Movie S1** (separate file to be uploaded separately).

File: MovieS1-DHAP-OH.mp4

Caption: Movie of ADMP trajectory of a hydroxide ion abstracting the *pro-R* proton (atom 22) from DHAP in the presence of Glu165. Route card = # b3lyp/6-31g(d,p) admp=(maxpoints=2000,nke=37052)

**Movie S2** (separate file to be uploaded separately)**.**

File: MovieS2.mp4

Caption: MP4 movie of Gaussian 09 energy minimization computation of hydroxide during barrierless abstraction of pro-R proton from dihydroxyacetone in presence of Glu165. Route= #opt b3lyp/6-31g(d,p)

**Movie S3** (separate file to be uploaded separately).

File: MovieS3.mp4

Caption: MP4 movie of Gaussian 09 energy minimization computation of a gas phase water dimer in an applied constant electric field of 0.065 au. Route = # opt b3lyp/cc-pVTZ nosym field=x-650 scf=(xqc,maxconventionalcycles=512
